# Reliable CRISPR/Cas9 genome engineering in *Caenorhabiditis elegans* using a single efficient sgRNA and an easily selectable phenotype

**DOI:** 10.1101/105718

**Authors:** Sonia El Mouridi, Claire Lecroisey, Philippe Tardy, Marine Mercier, Alice Leclercq-Blondel, Nora Zariohi, Thomas Boulin

**Affiliations:** Institut NeuroMyoGène Univ Lyon, Université Claude Bernard Lyon 1 CNRS UMR 5310, INSERM U1217 8 Rue Raphaël Dubois 69100, Villeurbanne, France

**Keywords:** CRISPR/Cas9 genome engineering, *C. elegans*, mScarlet, *dpy-10*, co-conversion

## Abstract

CRISPR/Cas9 genome engineering strategies allow the directed modification of the *C. elegans* genome to introduce point mutations, generate knock-out mutants and insert coding sequences for epitope or fluorescent tags. Three practical aspects however complicate such experiments. First, the efficiency and specificity of single-guide RNAs (sgRNA) cannot be reliably predicted. Second, the detection of animals carrying genome edits can be challenging in the absence of clearly visible or selectable phenotypes. Third, the sgRNA target site must be inactivated after editing to avoid further double-strand break events. We describe here a strategy that addresses these complications by transplanting the protospacer of a highly efficient sgRNA into a gene of interest to render it amenable to genome engineering. This sgRNA targeting the *dpy-10* gene generates genome edits at comparatively high frequency. We demonstrate that the transplanted protospacer is cleaved at the same time as the *dpy-10* gene. Our strategy generates scarless genome edits because it no longer requires the introduction of mutations in endogenous sgRNA target sites. Modified progeny can be easily identified in the F1 generation, which drastically reduces the number of animals to be tested by PCR or phenotypic analysis. Using this strategy, we reliably generated precise deletion mutants, transcriptional reporters, and translational fusions with epitope tags and fluorescent reporter genes. In particular, we report here the first use of the new red fluorescent protein mScarlet in a multicellular organism. wrmScarlet, a *C. elegans*-optimized version, dramatically surpassed TagRFP-T by showing an 8-fold increase in fluorescence in a direct comparison.

## Introduction

The pace of technical developments allowing the direct manipulation of genome sequences has seen a marked acceleration in the last years with the emergence of RNA-targeted nucleases derived from bacterial immune systems (Doudna and Charpentier 2014; Zetsche et al. 2015). In particular, the binary system relying on the *Streptococcus pyogenes* Cas9 endonuclease targeted by CRISPR (clustered, regularly interspaced, short, palindromic repeat) RNAs has been successfully used to generate point mutations, deletion or DNA insertions in an ever-growing number of experimental systems. *S. pyogenes* CRISPR/Cas9 has been adapted early on in the model nematode *C. elegans* (Friedland et al. 2013; Dickinson et al. 2013; Chen et al. 2013; Frøkjær-Jensen 2013; Dickinson and Goldstein 2016). Previously, heritable genome engineering could only be achieved in *C. elegans* – with significant effort – by remobilizing a Drosophila *Mos1* transposon, which could be inserted and excised in the germline (Robert and Bessereau 2007; Frøkjær-Jensen et al. 2010).

Despite great promise and early success, day-to-day CRISPR experiments are often not straightforward. Different factors might explain variability and inefficiency of CRISPR/Cas9 genome engineering in *C. elegans*. One specific reason could be the limited expression of heterologous genes in the germline due to dedicated co-suppression mechanisms (Kelly and Fire 1998). One approach to circumvent this problem has been to inject preassembled ribonucleoprotein (RNP) complexes of SpCas9 and CRISPR RNAs (crRNA – tracrRNA duplexes) directly into the germline(Cho et al. 2013; Paix et al. 2015). However, this approach is generally more expensive and less practical than using DNA expression vectors.

Another general reason for CRISPR failure is that efficacy and specificity vary tremendously between different single guide RNAs (sgRNA). Systematic analyses in different systems have led to the prediction that protospacers terminating by a single guanosine (GNGG) or ideally a double guanosine motif (GGNGG) are generally more effective (Doench et al. 2014; Farboud and Meyer 2015). To estimate the prevalence of such sites, we selected a set of 22 genes coding for two-pore domain potassium channel subunits and collected the sequences of all sgRNA target sites in and close to exons of these genes. On average, these 22 loci contained 138 ± 40 protospacers. We found that 20 ± 5% of these matched the GNGG motif, and only 5 ± 2% matched the GGNGG motif (Supplementary Table 1). Since, the proximity of an sgRNA to the target site has a positive impact on the likelihood to generate gene edits (Paix et al. 2014), it is therefore likely that few or no high efficiency sgRNAs will be situated close to a given target region.

One approach to compensate for low CRISPR/Cas9 activity has been to use selection strategies to increase the number of tested progeny. Antibiotic and phenotypic selection protocols have been adapted in *C. elegans* (Ward 2015; Dickinson et al. 2015; Norris et al. 2015; Dickinson and Goldstein 2016; Schwartz and Jorgensen 2016). They have the further advantage of reducing hands-on time and facilitate the detection of successful genome editing events. When phenotypic or antibiotic selection is not applicable, Co-CRISPR strategies can be used to increase the likelihood of identifying individuals with genome edits. These co-conversion approaches consist in injecting the sgRNA targeting a locus of interest together with a second sgRNA that targets a “marker gene” (Kim et al. 2014; Arribere et al. 2014). Progeny that carries a modification in the “marker” locus are then more likely to carry edits in the locus of interest. However, since two distinct sgRNAs do not necessarily cut with the same efficiency or in the same germ cell, effectiveness of traditional Co-CRISPR co-conversion is variable and mostly indicates a successful injection and expression of Cas9 and sgRNA.

While these major efforts have improved the efficiency of genome engineering in *C. elegans*, it is still not at a level where it can be considered to work routinely and easily in most labs. In addition, all available strategies require the protospacer sequence to be disrupted once the edit is generated to prevent further CRISPR/Cas9 cutting/activity. This almost always requires the introduction of point mutations in the protospacer adjacent motif (PAM) or in multiple bases of the protospacer. The consequences of such mutations in introns and up- or downstream regulatory regions are difficult, if not impossible to predict. Similarly, silent mutations in exons can have unfavorable effects due to codon usage bias. Therefore, it would be ideal if genome edits, in particular insertions or point mutations, could be generated without modification of the surrounding original genomic sequence.

Finally, since CRISPR/Cas9 guide RNAs are short 19-20 bp long sequences, there are often multiple closely matching sites (i.e. differing only by a few base pairs) in the genome that could be targeted, albeit at lower frequency. While algorithms have been developed to easily predict such potential off-target sites (Hsu et al. 2013; Doench et al. 2016), the prevalence of undesired CRISPR events has not been systematically analyzed in *C. elegans* and would require *ad hoc* experiments for each sgRNA.

We describe here a two-step strategy for reliable and scarless modification of the *C. elegans* genome using a single guide RNA that facilitates the detection of genome engineering events based on an easily selectable phenotype. Indeed, we reasoned that it should be possible to circumvent many practical hurdles described above if we transplanted the protospacer for a highly-efficient sgRNA into a genomic locus of interest to create an “entry strain” that would be more amenable to genome engineering. Specifically, we inserted a protospacer and PAM from the *dpy-10* gene(Arribere et al. 2014) – further referred to as the “*d10* site” or “*d10* sequence” – close to the targeted genomic region. In this “*d10*-entry strain”, we could then induce double-strand breaks at both the transplanted *d10* site and the endogenous *dpy-10* locus using a single sgRNA. We demonstrated that the *d10* site and the *dpy-10* locus were efficiently cut within the same nucleus. Finally, we found that co-conversion events (insertions of fluorescent reporter genes and epitope tags) occurred on average in 8 % (i.e. 1 in 12 animals) of F1 progeny that also carried mutations in the marker gene *dpy-10*, revealing a high incidence of co-conversion events. Since this co-conversion step no longer relied on an endogenous protospacer from the targeted locus, we did not need to introduce mutations in PAM or protospacer sequences and could generate perfectly accurate and scarless genome edits. Although our strategy is especially suited to insert sequences into the genome, we could also obtain large, precisely targeted gene deletions.

## Materials and Methods

### Strains generated in this study

N2 Bristol was used as a wild-type starting strain for transgenic lines generated in this study. Worms were raised at 20°C on nematode growth medium and fed *Escherichia coli OP50*. Worms were grown at 25°C after injection. Supplementary Table 2 provides a comprehensive list of the strains constructed for this study.

### Molecular Biology

Single guide RNA expression vectors (see Supplementary Methods) and plasmid repair templates were constructed using standard molecular biology techniques and Gibson assembly (Gibson 2011). They were systematically validated by Sanger sequencing before injection. Supplementary Table 3 and 4 respectively list the oligonucleotides and vectors used in this study. The Cas9-expression vector pDD162 was obtained from Addgene (Dickinson et al. 2013). Vectors generated for this study are available upon request.

### DNA preparation and microinjection

The pDD162, pMD8 and pPT53 plasmids were purified using the Qiagen EndoFree Plasmid Mega Kit (Qiagen). All other vectors were prepared using Invitrogen PureLink™ HQ Mini Plasmid Purification Kit (ThermoFisher Scientific). Single-strand DNA repair templates were synthetized and PAGE-purified by Integrated DNA Technologies (IDT). Except specified otherwise, plasmid vectors and ssDNA were diluted in water and injected at a final concentration of 50 ng/μL; co-injection markers were injected at 5 ng/μL. DNA mixes were injected into a single gonad of one day-old adult hermaphrodites raised at 20°C. They were then cloned onto individual plates after overnight incubation at 25°C.

### PCR screening

PCR DNA amplification was performed on crude worm extracts. In brief, worms were collected in ice-cold 1X M9 buffer, and 5 μL of packed worms were lysed by freeze thaw lysis in 14 μL of Worm Lysis Buffer (50 mM KCl, 10 mM Tris-HCl (pH = 8.3), 2.5 mM MgCl_2_, 0.45% Nonidet P-40, 0.45% Tween 20, 0.01% (w/v) Gelatin), to which 1 μL of proteinase K was added (1 mg/mL final concentration). After freezing at - 80°C, lysates were incubated for 1 hour at 65°C, and proteinase K was inactivated by further incubation at 95°C for 20 minutes.

High-fidely DNA polymerases (Q5^®^ High-Fidelity DNA Polymerase, New England Biolabs; Phusion High-Fidelity PCR Kit, Thermo Fisher Scientific) were used for PCR amplification to maximize the chances of recovery of desired modifications. Indeed, when we generated the *TagRFP-T::twk-18* knock-in strain, we initially screened 77 F1 clones using a low fidelity DNA Polymerase (Taq’Ozyme, Ozyme) and found no edits. When we immediately rescreened the same worm lysates with a more processive, high-fidelity DNA polymerase (Phusion, ThermoFisher Scientific) we identified 5 positive clones. PCR primers used for this study are listed in Supplementary Table 3.

### Generation of sgRNA expression vector by single strand DNA isothermal ligation

All sgRNA expression vectors were built using the novel pPT2 vector (see below and Supplementary Methods). In brief, pPT2 was linearized by PmeI/SexAI double digestion. The protospacer sequence was then inserted by isothermal ligation using a single-strand oligonucleotide containing the protospacer sequence flanked by 20 bp-long homology arms corresponding to the sequences upstream of PmeI and downstream of SexAI. If it was not already present in the sequence, a guanine residue was manually added 5’ to the protospacer sequence to optimize U6 promoter activity. A unique identifier was given to each sgRNA using the following nomenclature: CRpXYn, where “CR” stands for “CRISPR/Cas9 recognition site” and pXYn is the name of the corresponding sgRNA-expression plasmid.

### Codon-optimization of mScarlet

wrmScarlet was generated by gene synthesis (Gblock, IDT) based on the mScarlet sequence (Genbank KY021423; (Bindels et al. 2017); Figure 3 Supplement 1). Codon-optimization was performed using the “*C. elegans* codon adapter” service (Redemann et al. 2011) with the following parameters: “*0 introns*”, “*optimize for weak mRNA structure at ribosome binding site”,* and “*avoid splice sites in coding region*”. The Gblock fragment library was combined by isothermal ligation with left and right homology regions flanking the *d10* sequence in *twk-18(bln213)* to generate the repair template pSEM87. The wrmScarlet cDNA sequence is available upon request.

### Microscopy and fluorescence quantification

Freely-moving worms were observed on NGM plates using an AZ100 macroscope (Nikon) equipped with a Flash 4.0 CMOS camera (Hamamatsu Photonics). Confocal imaging was performed using a Nikon Eclipse Ti inverted microscope equipped with a CSUX1-A1 spinning-disk scan head (Yokogawa) and an Evolve EMCCD camera (Photometrics). Worms were imaged on 2% fresh agar pads mounted in M9 solution containing 50 mM NaN_3_.

Comparison of wrmScarlet and TagRFP-T fluorescence was performed as follows: (1) confocal stacks of the head region were acquired for TagRFP-T and wrmScarlet knock-in strains on the same day, using identical settings, and NaN_3_ immobilization; (2) the same number of confocal slices was selected from each stack; (3) stacks were projected by summing fluorescence at each pixel position in the stack; (4) total fluorescence was measured in areas of identical size and position relative to the anterior tip of the worm and pharynx; (5) total fluorescence was corrected by subtracting equipment noise, i.e. fluorescence measured in an area of the same size outside of the sample.

### Data and reagent availability

All *C elegans* strains and plasmids described in this study are available upon request.

## Results

### Generation of *d10*-entry strains as a starting point for robust and precise gene modification

The starting point of our strategy consists in the insertion of the *d10* sequence (i.e. *dpy-10* protospacer + PAM) into the locus of interest (Figure 1A). First, we targeted three positions in two genes coding for two-pore domain potassium channel subunits: (1) the ATG start site of *sup-9*, (2) the ATG of the *egl-23b* isoform and (3) the common stop codon of all *egl-23* isoforms (Figure 1A). Next, we predicted all possible sgRNA sequences within a 50-base window around these positions, and selected sgRNAs close to the ATG or stop codons. Using multiple sgRNAs increases the chances of finding at least one sgRNA that cuts efficiently enough to insert the *d10* site at the desired location. We then defined the portion of the gene to be replaced by the *d10* site, based on the positions of the most upstream and most downstream PAM sequences. Finally, we designed a single-strand oligonucleotide sequence (ssON) containing the *d10* sequence flanked by up- and downstream homology regions of approximately 50 bases (Figure 1A). This ssON could serve as a repair template with all selected sgRNAs since it did not contain their protospacer or PAM sequences.

**Figure 1.**
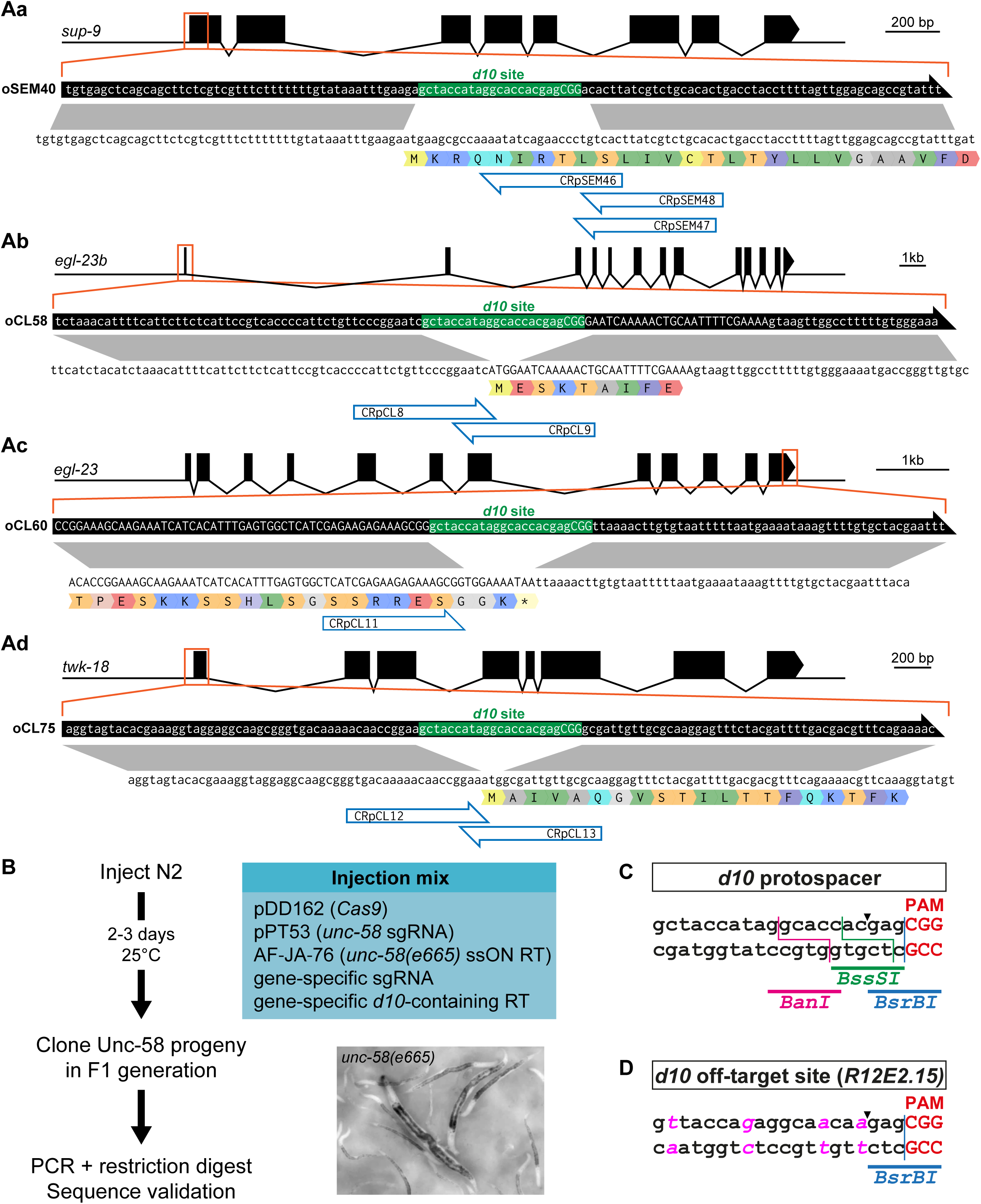
Generation of *d10*-entry strains. **A** Insertion of the *d10* sequence into **(Aa)** *egl-23b,* **(Ab)** *egl-23 C-term,* **(Ac)** *sup-9*, and **(Ad)** *twk-18* using a single-strand oligonucleotide repair template compatible with multiple sgRNAs. Genes and their intron/exon structure are displayed in the 5’ to 3’ orientation. The ssON repair templates are represented by black arrows (containing the *d10* sequence in green) above the coding strand and translation of the target gene. Correspondence of homology regions between the ssON repair template and genomic locus is indicated in gray. sgRNA binding sites are indicated by blue open arrows. **B** *unc-58* co-conversion is used to detect the insertion of *d10* sequences into a geneof interest. *unc-58(e665)* mutants are easily identified in the F1 progeny of injected P0 animals based on their straight body posture, lack of mobility and characteristic rotation around the antero-posterior body axis. RT, repair template. **C** BanI, BssSI, and BsrBI restriction sites are present in the *d10* protospacer sequence and are used for RFLP analysis. The Cas9 double-strand break site is indicated by an arrowhead. **D** *R12E2.15* contains the only predicted off-target site of the *d10* sgRNA. Four base changes (in pink) distinguish both sites. A BsrBI site follows the Cas9 double-strand break site (indicated by an arrowhead), between the -3 and -4 bases relative to the protospacer adjacent motif (PAM).

Next, we built the necessary sgRNA expression constructs using a novel vector and assembly strategy. This vector (pPT2) is composed of an RNA Polymerase III U6 promoter from *K09B11.12* (Friedland et al. 2013; Katic et al. 2015) followed by two restriction sites (PmeI and SexAI), followed by the sgRNA portion corresponding to the CRISPR tracrRNA and 3’ UTR of *K09B11.12*. This vector was linearized by restriction digest with PmeI and SexAI, and the crRNA sequence was incorporated by isothermal ligation (Gibson assembly (Gibson 2011)) using a single single-strand DNA oligonucleotide (see Materials and Methods and Supplementary Methods). These sgRNA expression vectors were systematically validated using Sanger sequencing.

Since it is not possible to predict the efficiency of an sgRNA *a priori*, we reasoned that we could increase the likelihood of finding a *d10* insertion at the locus of interest by using a moderately efficient Co-CRISPR. We chose a previously described reagent combination that introduces a mutation in the two-pore domain potassium channel *unc-58* and replicates the L428F amino acid change found in the *unc-58(e665)* reference allele (Arribere et al. 2014). *unc-58(e665)* produces a dominant and easily detectable phenotype. Worms have a straight body posture and are essentially unable to move on solid NGM medium throughout their post-embryonic development (Figure 1B). However, they are viable and fertile. *unc-58(e665)*-like progeny can be detected two to three days post-injection and individual F1 worms can be cloned right away to ensure that independent events are selected.

To generate *d10-entry* strains for *sup-9* and *egl-23*, we injected wild-type N2 worms with a mix of plasmid DNA and ssON repair templates (Figure 1B). In each case the mix was composed of (i) a Cas9 expression vector (pDD162), (ii) the sgRNA expression vector targeting *unc-58* (pPT53), (iii) one gene-specific sgRNA expression vector, (iv) the ssON to introduce the *e665* mutation AF-JA-76(Arribere et al. 2014), and (v) the ssON required to introduce the *d10* site (Figure 1A, Supplementary Table 3). After three to four days, we cloned Unc-58-marked F1 worms to single plates. We then detected the presence of the *d10* site in the F2 population by PCR amplification and restriction digest. The *d10* sequence contains sites for three restriction enzymes (BanI, BsrBI, and BssSI) that can be used for restriction fragment length polymorphism analysis (RFLP) (Figure 1C). In each case, we designed a PCR primer pair that produced a fragment of 500 to 600 bp, centered on the *d10* site. In this way, we were able to generate multiple independent *d10*-entry strains for each of the targeted loci (Table 1 and Supplementary Table 2). In each case, we selected homozygous clones for the *d10* insertion that lacked the *unc-58* gain-of-function mutation, and validated them by Sanger sequencing around the *d10* sites.

**Table 1:**
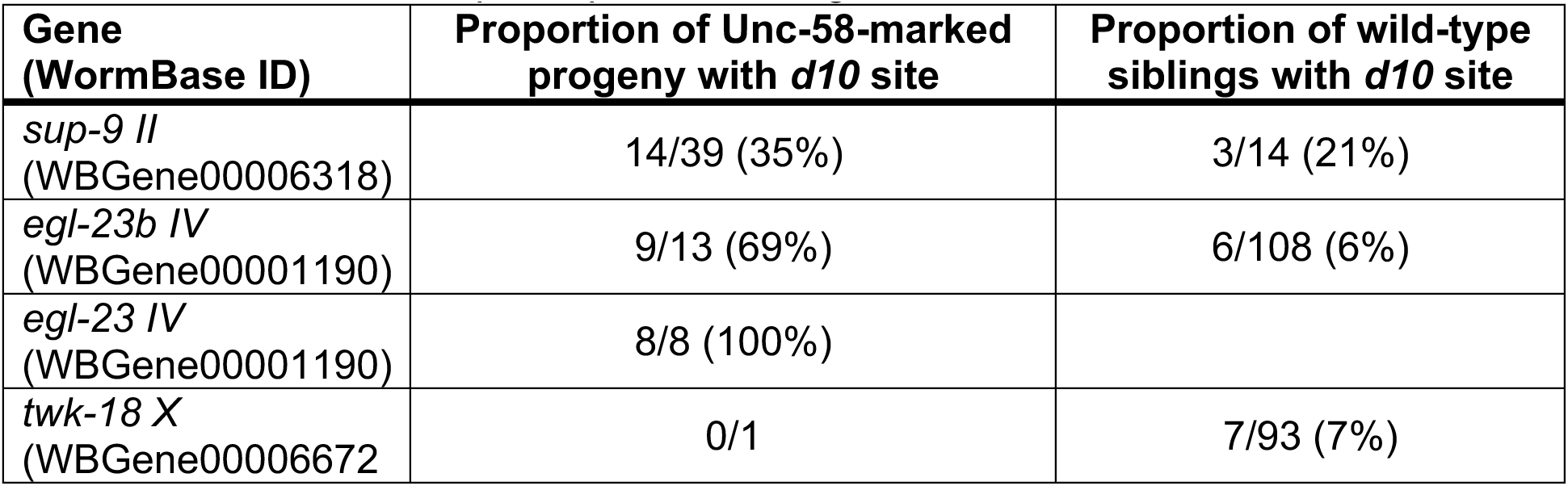
Insertion of *d10* protospacer at four genomic loci.

Next we targeted the two-pore domain potassium channel subunit *twk-18* (Figure 1Ad). In this experiment, only one of 41 injected P0 worms gave a single Unc-58 worm (Table 1). Since this marked F1 worm did not incorporate the *d10* site in *twk-18*, we decided to screen its unmarked siblings. Doing so, we found 7 independent insertion events out of 93 tested clones. Similarly, we found 3 additional *d10* insertion events in 14 unmarked siblings of the *sup-9* experiment, and 6 additional *d10* insertions in 108 unmarked siblings of the experiment targeting the ATG of the *egl-23b* splice variant (Table 1).

In conclusion, screening for Unc-58-marked F1 progeny allowed us to rapidly identify P0 individuals for which the injection was successful and CRISPR/Cas9 activity was present in the germline. Cloning Unc-58 worms at the F1 generation ensured that we selected independent edits and decreased the number of animals to clone and analyze by PCR. In three cases, we also found *d10* protospacer insertions in non-marked siblings, although at lower frequencies than in Unc-58-marked F1 progeny. In total we successfully targeted 5 different sites in the genome using this protocol (Supplementary Table 2).

### Efficient and specific cutting of transplanted *d10* sites

Different laboratories have independently reported that the sgRNA targeting the *d10* site is among the most efficient ones currently known (I. Katic, M. Boxem, C. Gally, J.-L. Bessereau, personal communication). The reasons for this high efficacy are unclear. For example, the site matches the GNGG motif and not GGNGG (Farboud and Meyer 2015). A more favorable chromatin organization or the sequence of the *dpy-10* locus itself might explain high CRISPR activity in this gene. Since we transplanted only the protospacer and PAM sequences of the *d10* site, we decided to estimate the frequency of cuts in transplanted *d10* sites before attempting to engineer these loci by homologous recombination.

DNA double-strand breaks can be repaired by homologous recombination using the sister chromatid to restore a wild-type sequence or by non-homologous end joining (NHEJ), which results in small indels close to the cut site. We reasoned that we could therefore estimate the double-strand break frequency by looking for the destruction of the restriction sites present in and around the -3/-4 position relative to the NGG, i.e. the Cas9 cut site (Figure 1C). Note that only catastrophic events that result in sufficiently modified *d10* sites that could no longer be targeted by the Cas9/*d10*-sgRNA duplex would be detected in this way. This experiment therefore underestimates the double-strand break frequency since precise repair events using the sister chromatid would not be detected.

We selected four *d10*-entry strains on three different chromosomes (*tag-68 I*, *egl-23 IV*, *twk-18 X* and *unc-58 X*). Each strain was injected with a DNA mixture containing (i) a Cas9 expression vector (pDD162), (ii) an sgRNA expression vector targeting *dpy-10* (pMD8), and (iii) a ssON to introduce the *cn64* mutation (AF-ZF-827) in *dpy-10*(Arribere et al. 2014). Next, we singled F1 progeny showing a Dpy-10 phenotype, i.e. Rol (*cn64*/+), Dpy (-/-) or DpyRol (*cn64*/-)(Levy et al. 1993; Arribere et al. 2014). Finally, we tested all clones that segregated the Dpy-10 phenotype in their progeny and observed the loss of the BanI site in 14 to 26 % of them (Table 2). Since BanI is located 5’ to the cut site (Figure 1C), we tested the remaining BanI-positive clones (i.e. lacking mutations in BanI) with BsrBI and BssSI. This lead us to identify additional events, likely affecting the bases closest to the -3/-4 cut site. In total, we found that between 33 and 52% of Dpy-10-marked F1 worms had lost at least one restriction site, which demonstrates that heterologous *d10* sites can be cut at high frequency and are present in Co-CRISPR-marked F1 progeny.

**Table 2:**
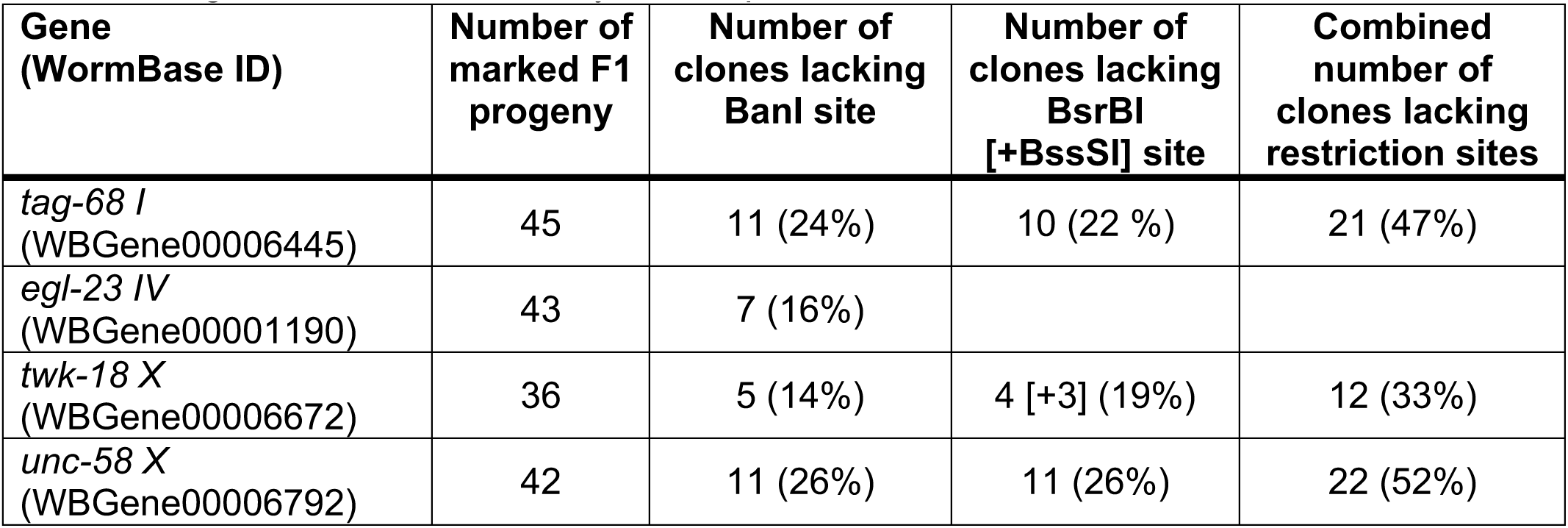
High CRISPR/Cas9 activity at transplanted *d10* site.

Bioinformatic analysis predicts a single, low scoring, off-target site for the *d10* sgRNA, situated in the uncharacterized gene *R12E2.15* (Figure 1D). We investigated potential off-target cutting of the *d10* sgRNA by analyzing the *R12E2.15* locus in 32 independent F1 worms that segregated the Dpy-10 phenotypes. None of these 32 lines showed scars around the potential off-target cut site of the *d10* sgRNA.

Given the high correlation between worms displaying Dpy-10 phenotypes and double-strand break events in the transplanted *d10*-site, and given the high selectivity of the *d10* sgRNA for the endogenous and transplanted sites, we chose to focus only on Dpy-10-marked Co-CRISPR individuals in our co-conversion experiments.

### Generation of multiple knock-in lines using a single *d10*-entry strain

As a proof of principle for our strategy, we targeted the *twk-18* locus. TWK-18 is one of 47 two-pore domain potassium channels in the *C. elegans* genome. Its expression pattern and localization in body wall muscle cells has been reported previously (Kunkel et al. 2000). We decided to generate two N-terminal fusions (1) with the red fluorescent protein TagRFP-T (Shaner et al. 2008) and (2) with the blue fluorescent protein TagBFP (Chai et al. 2012). As a repair template, we constructed two vectors with left and right homology regions of 2073 and 1993 base pairs (Figure 2A). We injected each repair template separately into the *twk-18 d10*-entry strain (JIP1143) with (i) a Cas9 expression vector (pDD162), (ii) the sgRNA expression vector targeting *dpy-10* (pMD8), (iii) the ssON to introduce the *cn64* mutation in *dpy-10* (AF-ZF-827) and (iv) the fluorescent reporter pCFJ90 as a co-injection marker to identify transgenic animals based on mCherry fluorescence in the pharynx (Figure 2B). We selected 77 (TagBFP) and 98 (TagRFP-T) Dpy-10-marked F1 progeny. Finally, we used PCR screening to identify 5 and 6 clones respectively, which had integrated the TagRFP-T and TagBFP sequences in the *twk-18* locus, corresponding to a recombination frequency of 6% of Dpy-10-marked F1 progeny (Table 3).

**Figure 2.**
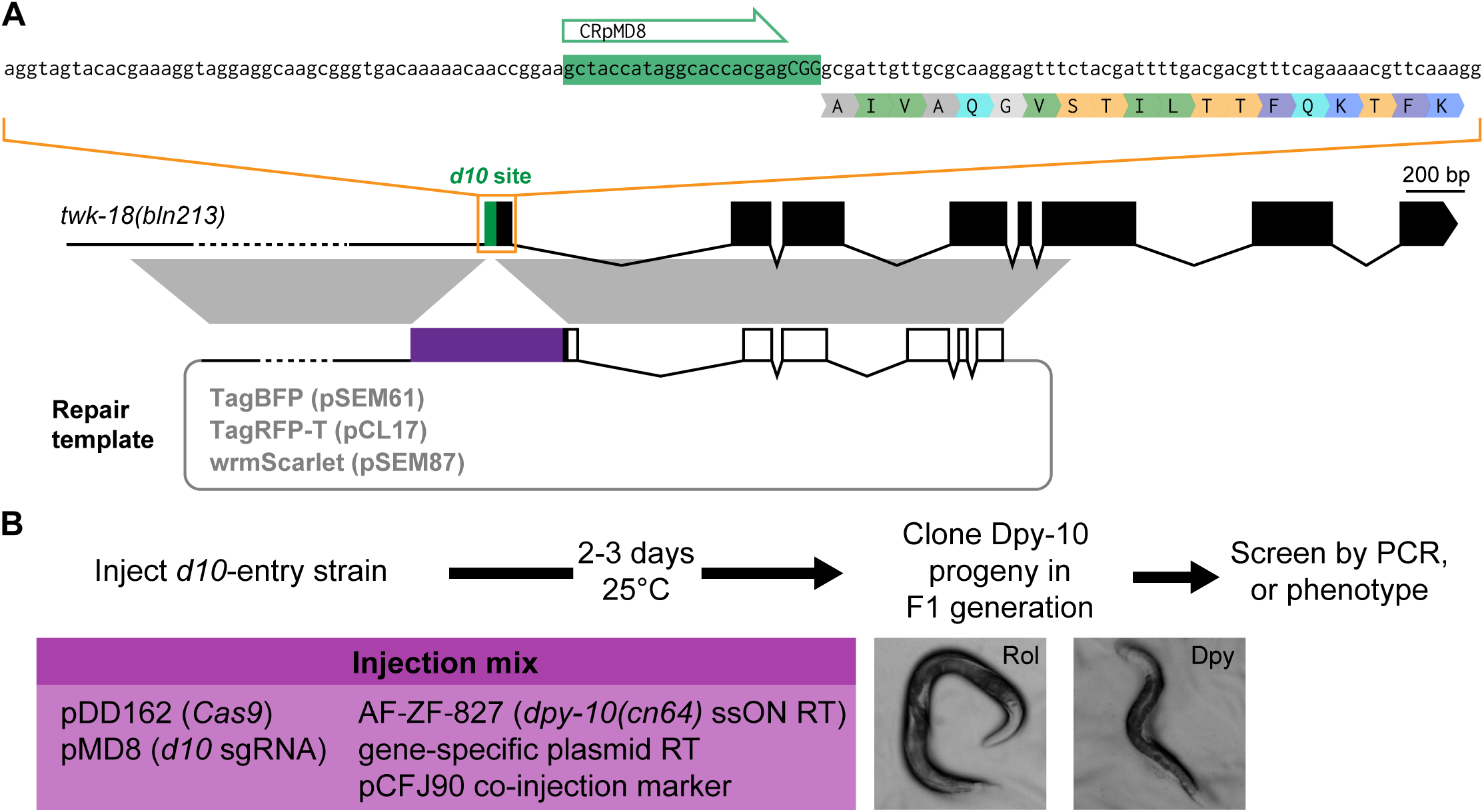
Generation of multiple knock-in lines using a single *d10*-entry strain. **A** single *d10-*entry strain is used to engineer N-terminal TagBFP, TagRFP-T, and wrmScarlet fusions in the *twk-18* locus. Correspondence of homology regions between the plasmid repair template and *twk-18* genomic locus is indicated in gray. RT, repair template. **B** Two to three days following injection of a *d10*-entry strain with a CRISPR/Cas9 mix, F1 progeny with Dpy-10 phenotypes (Rol or Dpy) can be easily recovered, and further screened in the F2 generation to identify the desired genome edits by PCR or phenotype.

**Table 3:**
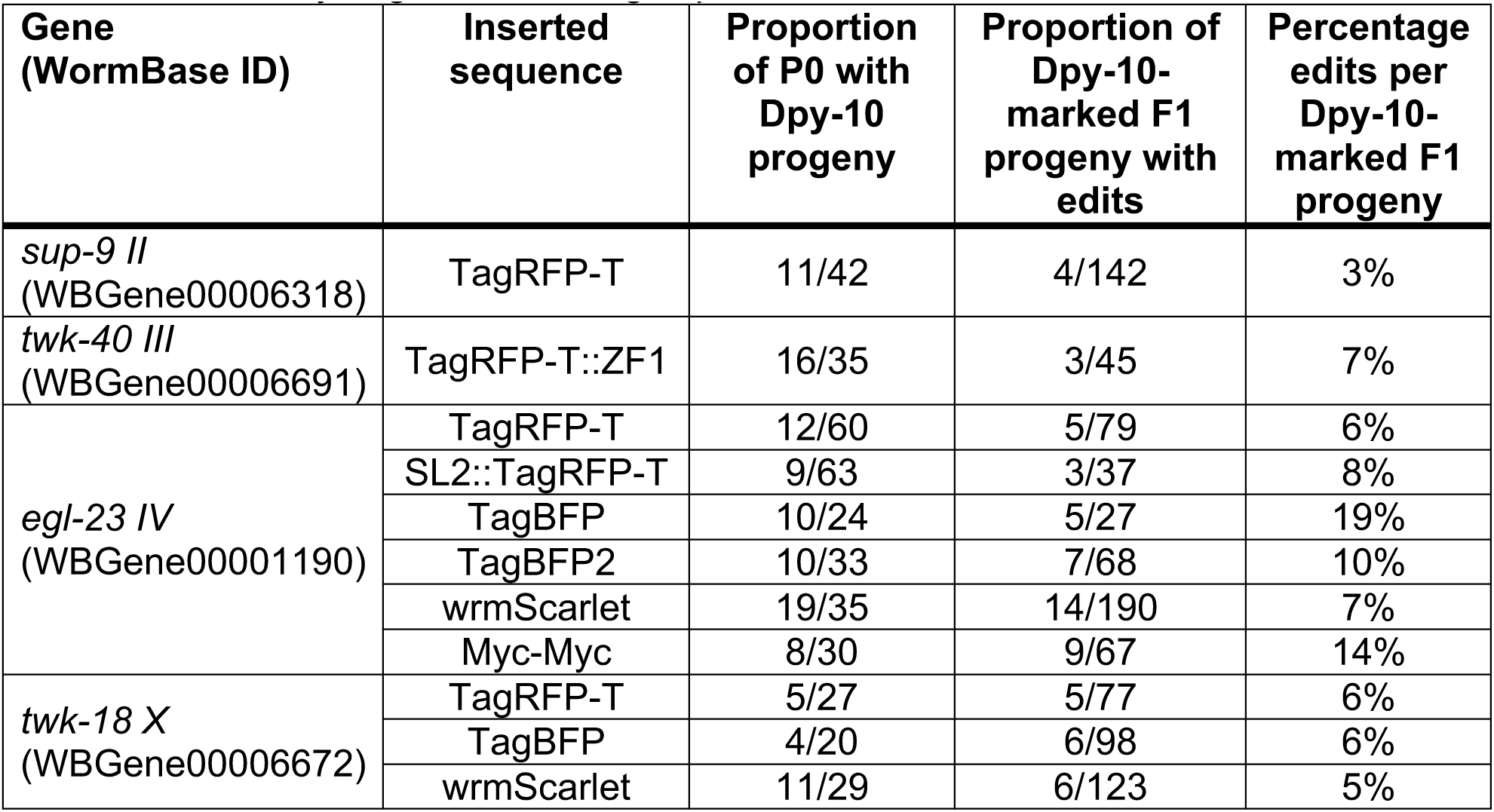
Summary of genome editing experiments.

When we prepared these knock-in lines for observation by confocal fluorescence microscopy, we noted that TWK-18-TagBFP had a very reproducible subcellular distribution at the exterior surface of body wall muscle cells (Figure 3B). The highly repetitive grid-like pattern was very different from the one reported previously since it appeared to show strong GFP signal in the endoplasmic reticulum (Kunkel et al. 2000). This intracellular localization was not consistent with the electrophysiological effect of TWK-18 gain-of-function mutants, in which TWK-18 most likely exerts its hyperpolarizing role at the plasma membrane. We believe these differences probably resulted from a strong over-expression of TWK-18 in this study compared to our knock-in strain, highlighting the importance of physiological expression levels when observing the distribution of cell surface-targeted channels and receptors (Gendrel et al. 2009). When comparing the TagRFP-T and TagBFP knock-in strains, we noticed a marked difference in brightness but also in the apparent resolution (Figure 3B). The overall pattern of TagRFP-T was similar to TagBFP but the longer emission wavelength of TagRFP-T (emission maximum, 584 nm) did not afford the same resolution as the much shorter emission wavelength of TagBFP (emission maximum, 457 nm). This is in part explained by the fact that resolution is proportional to the emission wavelength, making TagBFP an interesting alternative to increase imaging resolution without changing imaging hardware.

**Figure 3.**
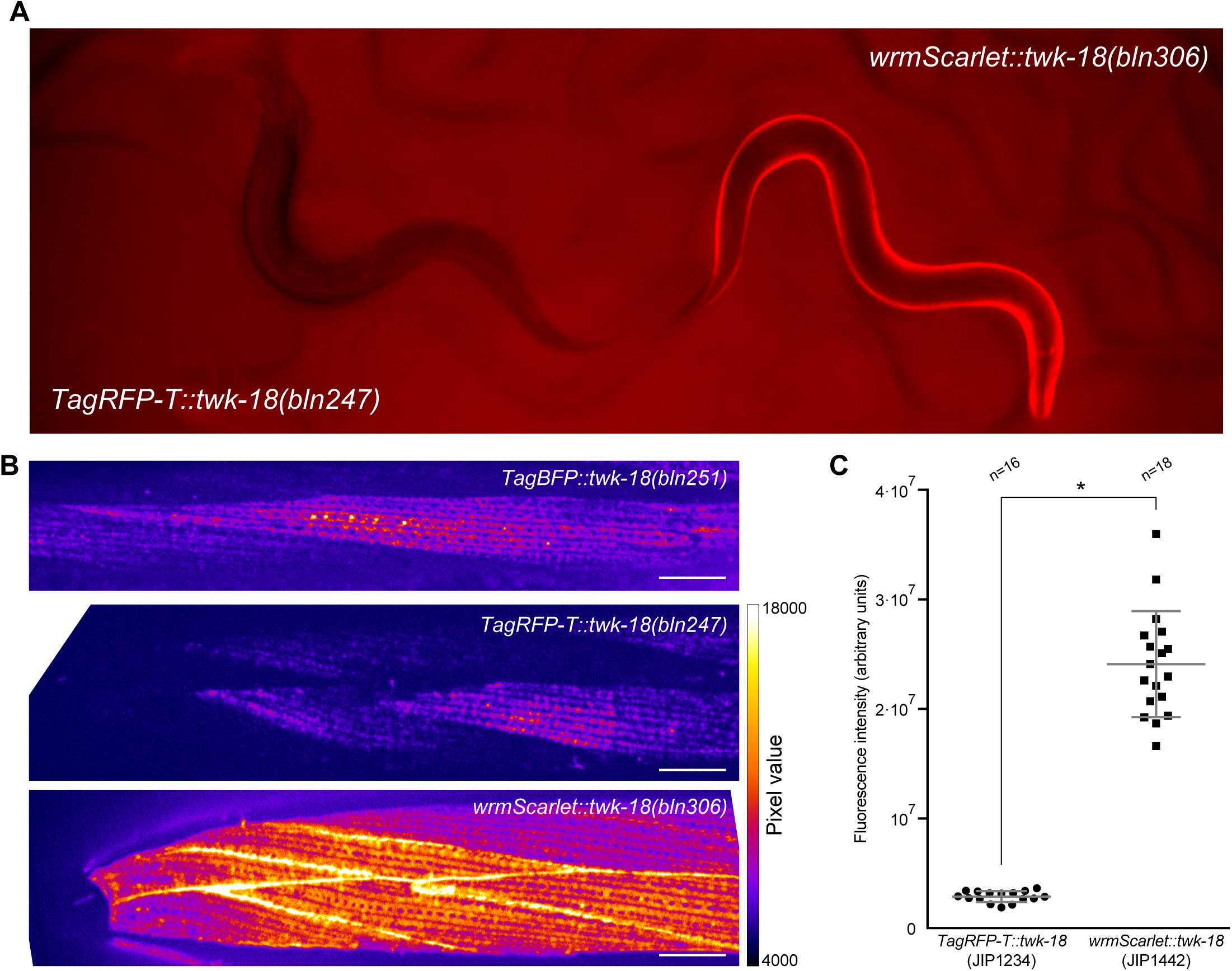
Comparison of TagBFP, TagRFP-T and wrmScarlet using reliable editing of the *twk-18 locus*. **A** wrmScarlet::TWK-18 is visibly brighter than TagRFP-T::TWK-18. Side-by-side comparison of two young adult hermaphrodites. wrmScarlet-associated fluorescence is visible by eye in freely moving worms on NGM plates, while TagRFP-T is not detectable by eye in this context. **B** The two-pore domain potassium channel TWK-18 decorates the plasma membrane of body wall muscle cells. Representative images of head muscle cells labeled with N-terminal fusions of TWK-18 to TagBFP, TagRFP-T, and wrmScarlet. Head is left. Scale bars, 10 μm. **C** Quantification of fluorescence intensity shows an 8-fold increase in fluorescence between TagRFP-T and wrmScarlet. Mean ± standard deviation. Student’s *t*-test, * < 0.0001.

Next, we targeted three additional loci on different chromosomes (*sup-9 II*, *twk-40 III*, and *egl-23 IV*) and generated seven different edits with a variety of insert types (TagRFP-T, TagRFP-T::ZF1, SL2::TagRFP-T and TagBFP) (Table 3 and Supplementary Table 2). We found that we could reliably edit these different loci. Indeed, edit frequencies in Dpy-10-marked F1 worms ranged from 3% to 19% (average 8 %). Taken together, these experiments demonstrate that it is possible to take advantage of the high CRISPR activity of the *d10* sgRNA to robustly engineer the genome of *C. elegans*. This strategy significantly reduces hands-on work by focusing only on the animals that most likely carry genome edits. It generates scarless edits since it does not require the introduction of mutations in endogenous protospacer sequences.

### wrmScarlet, a brighter red fluorescent protein

The development of improved blue (TagBFP, (Chai et al. 2012)), cyan (mTurquoise2, (Goedhart et al. 2012)), green (mNeonGreen, (Shaner et al. 2013)), and red fluorescent proteins (TagRFP-T, (Shaner et al. 2008)) has greatly increased our capacity to detect proteins expressed at physiological levels. However, the properties of these new fluorophores are generally characterized in bacteria or cell culture systems, and are not always retained in *C. elegans* cells or in specific subcellular compartments (Heppert et al. 2016).

In an effort to improve the detection of fusion proteins *in vivo*, we have investigated the behavior of the recently described red fluorescent protein mScarlet (Bindels et al. 2017). mScarlet has currently the highest reported brightness, quantum yield and fluorescence lifetime of any red fluorescent protein. We synthetized a *C. elegans* codon-optimized cDNA (Redemann et al. 2011) of mScarlet, which we named wrmScarlet (Figure 3 Supplement 1). We combined this cDNA with homology arms flanking the *d10* site in *twk-18* to generate a *wrmScarlet*::*twk-18* repair plasmid (pSEM87, Figure 2A, and Figure 3 Supplement 1). Following the same strategy as before, we injected 29 P0 worms (*twk-18 d10*-entry strain, JIP1440) with an injection mix containing (i) pDD162 (Cas9), (ii) the ssON to introduce the *cn64* mutation in *dpy-10* (AF-ZF-827), (iii) the sgRNA expression vector targeting *dpy-10* (pMD8), and (iv) *wrmScarlet::twk-18* repair plasmid (pSEM87). Out of 29 injected P0 worms, 11 produced Dpy-10 F1 progeny. In total, we analyzed 123 Dpy-10-marked F1 worms that segregated Dpy-10 progeny and found 6 clones incorporating the wrmScarlet sequence (Table 3).

While undetectable by eye, specific fluorescence can be observed on NGM plates in *TagRFP-T::twk-18* worms with a macroscope (Nikon AZ100) coupled to a CMOS camera (Flash 4, Hamamatsu Photonics). Using the same macroscope, acquisition parameters and filter sets, wrmScarlet-TWK-18 was significantly brighter than the TagRFP-T fusion, so much so that it became visible to the naked eye (Figure 3A). We next compared the subcellular distribution and brightness of these two translational fusions using spinning disk confocal imaging. Both protein fusions had grossly identical distribution patterns (Figure 3B). However, the wrmScarlet fusion was approximately eight times brighter than the TagRFT-T fusion in this assay (Figure 3C). In fact, the distribution of the wrmScarlet::TWK-18 fusion protein appeared more uniform than TagRFP-T::TWK-18, possibly due to the increased fluorescent signal, which compensated for the reduced resolution when compared to TagBFP (Figure 3B). These properties make wrmScarlet a very convincing replacement for TagRFP-T and should greatly facilitate the detection of protein fusions expressed at low, physiological expression levels.

### Generation of an epitope-tagged knock-in using a long single-strand oligonucleotide

For short edits, single-strand DNA oligonucleotides can be very efficient repair templates (Zhao et al. 2014; Arribere et al. 2014; Katic et al. 2015). We tested if a large ssON could be used as a repair template to integrate two repeats of the myc-tag sequence into the *egl-23* locus (Figure 2 Supplement 1). We synthetized a 182 nucleotide-long ssDNA fragment containing part of last exon of *egl-23* to restore the full-length C terminal sequence, followed by 75 nucleotides encoding two myc tag sequences, and the original stop codon and 3’ UTR region of the *egl-23* gene (Supplementary Table 3). In theory, each strand could serve as a template for recombination, but we selected the strand complementary to the sgRNA following the observations of (Katic et al. 2015). We injected 30 P0 worms (JIP1150) with a DNA mixture containing (i) a Cas9 expression vector (pDD162), (ii) the expression vector for the *d10* sgRNA (pMD8), (iii) the ssON that introduces the *cn64* mutation in *dpy-10* (AF-ZF-827), (iv) ssON containing the 2xMyc tag sequence (oSEM158) and (v) pCFJ90 as a co-injection marker to identify transgenic animals based on mCherry fluorescence in the pharynx. We selected 67 Dpy-10-marked F1 progeny and among these, 9 carried the 2xMyc tag. This 14% edit frequency was comparable, yet slightly higher than the average efficiency of longer inserts using double-strand DNA repair templates (Table 3).

The high edit efficiency observed in this experiment shows that our strategy is very effective to tag proteins of interest for immunohistochemical or protein biochemistry experiments. Generating this epitope-tagged strain required less than two weeks, with no additional cloning steps and could be repeated easily to integrate a variety of epitope tags, opening the way for different downstream applications.

### Generation of a large, targeted deletion using a *d10*-entry strain

One starting point for many CRISPR experiments is the desire to engineer loss-of-function mutations in a gene of interest. Previously, researchers relied on random mutagenesis with chemical mutagens or ionizing radiation followed by fastidious screening and extensive backcrossing to wild-type strains to eliminate background mutations (Boulin and Hobert 2012). CRISPR/Cas9 engineering offers the possibility to generate gene deletions with minimal background mutations. A simple approach relies on the repair of double-strand breaks by NHEJ pathways (Friedland et al. 2013; Chen et al. 2013; Katic and Großhans 2013; Waaijers et al. 2013; Dickinson and Goldstein 2016). One, two or more sgRNAs are injected together and phenotypic or PCR screening strategies are used to retrieve deletion mutants by PCR amplification. However, the exact breakpoints of these deletions are not controllable in this scheme and there is always the potential for undesired edits due to off-target effects for each sgRNA.

*d10*-entry strains can also serve as a starting point to generate precisely defined gene deletions. As a proof of principle, we targeted the *egl-23* locus. *egl-23* is a large locus comprising 12 exons, and removal of the entire *egl-23a* splice isoform required an 8 kb deletion (Figure 2 Supplement 1). Our goal was to replace the complete *egl-23*a locus by a transgene expressing the red fluorescent protein mCherry in the pharynx which could be used as a genetic balancer and knock-out mutant of *egl-23*. We therefore constructed a repair template composed of two homology regions of 2 kb, which flanked the transcriptional reporter unit (*Pmyo-2::mCherry::unc-543’UTR*). This construct was then injected into the appropriate *egl-23 d10*-entry strain (JIP1150) along with (i) a Cas9 expression vector (pDD162), (ii) the expression vector for the *d10* sgRNA (pMD8), (iii) the ssON that introduces the *cn64* mutation in *dpy-10* (AF-ZF-827). Out of 40 Dpy-10 progeny, we identified one knock-out line (JIP1253). We validated that the genome edit was accurate by Sanger sequencing. We further verified that the possible off-target site of the *d10* sgRNA was unaffected (Figure 1D). While this particular trial was less efficient than smaller insertions, it confirmed that *d10*-entry strains can – as expected – be used to generate large deletion and gene replacements, in addition to being ideally suited for the insertion of kilobase-sized inserts or epitope tags.

## Discussion

The major conceptual innovation of our strategy is to render genes highly susceptible to CRISPR/Cas9 engineering by transplanting the *d10* sequence. As we have described above, highly effective sgRNAs matching the GGNGG motif are underrepresented in the genome, and are therefore rarely found in close proximity to the region of interest. While editing frequency is highly variable between sgRNAs at different loci or even within the same locus, we found that editing using our *d10* strategy was robust at different loci, with edit frequencies averaging 8 %, i.e. 1 in 12 F1 progeny. In addition, editing was also robust at a single locus. This is particularly valuable and time-saving when multiple edits need to be generated in the same locus, as is usually the case when a gene is being characterized in depth. For example, we took advantage of this high editing frequency to rapidly generate multiple chromatic variants in the *twk-18* locus. This allowed us for the first time to precisely compare the resolution and fluorescence intensity of TagBFP, TagRFP-T and the recently published mScarlet. Based on the highly stereotypical distribution pattern of TWK-18 at the muscle surface, we could show that (1) TagBFP fusions provided the best apparent resolution, (2) a codon-optimized wrmScarlet was approximately 8 times brighter than TagRFP-T, (3) the increased signal of wrmScarlet partly compensated for the lesser resolution of red vs. blue fluorescent proteins.

One unique feature of our strategy is that edits can be designed so that all original genomic sequences are perfectly preserved. Indeed, by using the transplanted *d10* sgRNA instead of sgRNAs from the targeted locus, no mutations need to be introduced to avoid continued CRISPR/Cas9 activity once the edit is performed. This facilitates and accelerates experimental design, because only one repair template is designed instead of specific repair constructs for each endogenous sgRNA.

Another benefit of our strategy is that we could retrieve multiple independent lines from the same injected animal by cloning animals in the F1 generation, which is not possible in strategies that rely on the screening of mixed populations of F2 progeny, e.g. with antibiotic selection strategies. Therefore, since we could focus on relatively few F1 clones, multiple methods could be used to detect the desired genome edit such as direct observation, phenotypic screening or PCR detection. Limiting the number of animals that need to be analyzed, could also mitigate PCR detection issues (see Materials and Methods, PCR screening).

From a practical perspective, our strategy provides multiple layers of quality control. Based on the easily detectable Dpy-10 Co-CRISPR phenotype, we could directly monitor the success of injections and assess the general efficiency of the experiment over time and between experimenters. We could determine if an experiment would likely be successful within three days post-injection, by monitoring the number of marked F1 progeny. Finally, all steps of our protocol are only limited by the generation time of *C. elegans*, making it particularly time-efficient.

Obtaining the *d10*-entry strain is the major bottleneck of our strategy. This step, like every CRISPR/Cas9 experiment, relies (1) on the ability to find an endogenous sgRNA that cuts efficiently, and (2) on the rate of homology directed repair at the cut site, which could be influenced by the local genomic context or specific sequence features of the homology arms. Insertion of the *d10* sequence using single-strand oligonucleotides proved highly successful in most cases (Table 1). However, only one of the different sgRNAs we had selected gave us edits, highlighting again the variable efficacy of endogenous sgRNAs. During this study, we were unable to recover *d10* insertions in some of our target loci despite testing multiple sgRNAs. For some genes, we eventually succeeded by using double-stranded DNA repair templates with long homology arms instead of single-strand oligonucleotides. Another avenue we have begun to explore, is to use a CRISPR/Cas9 RNP complex instead of Cas9 expression plasmids. In one experiment, we were able to retrieve two *d10* insertions in this way, while injection of plasmids had been unsuccessful (data not shown).

Another practical concern appears when targeting loci that are closely linked to *unc-58* (to build *d10* entry strains) *or dpy-10* (to engineer *d10* loci), which are situated at the center of chromosome 2 and X, respectively. Since we select F1 progeny based on mutation of *dpy-10* or *unc-58*, it is likely that genome edits will be linked to these marker mutations. In that case, one should consider the wild-type siblings in the progeny of an injected P0 individual that produced a significant fraction of marked progeny.

So far we have tested this strategy only with the *d10* sequence, but in principle, any highly effective sgRNA that targets a gene producing a dominant Co-CRISPR phenotype could be used. Conceptually, our strategy could also be extended to other genetic model organisms. In particular, a co-CRISPR strategy based on the *white* locus has been recently published, and could be a starting point to adapt this strategy to engineer the Drosophila genome (Ge et al. 2016).

## Acknowledgements

We thank J.-L. Bessereau, I. Katic and members of the Bessereau team for helpful discussion, M. Jospin for comments on the manuscript, and Manuela D’Alessandro for constructing pMD8. pCFJ90 was a gift of Christian Frøkjær-Jensen. We thank Wormbase, which is supported by National Institutes of Health (NIH) grant U41 HG002223. This work was supported by a research grant from Fondation Fyssen (T. B.) and an ERC Starting Grant (Project *Kelegans*) (T. B.).

## Supplementary Figures

**Figure 2 Supplement 1.**
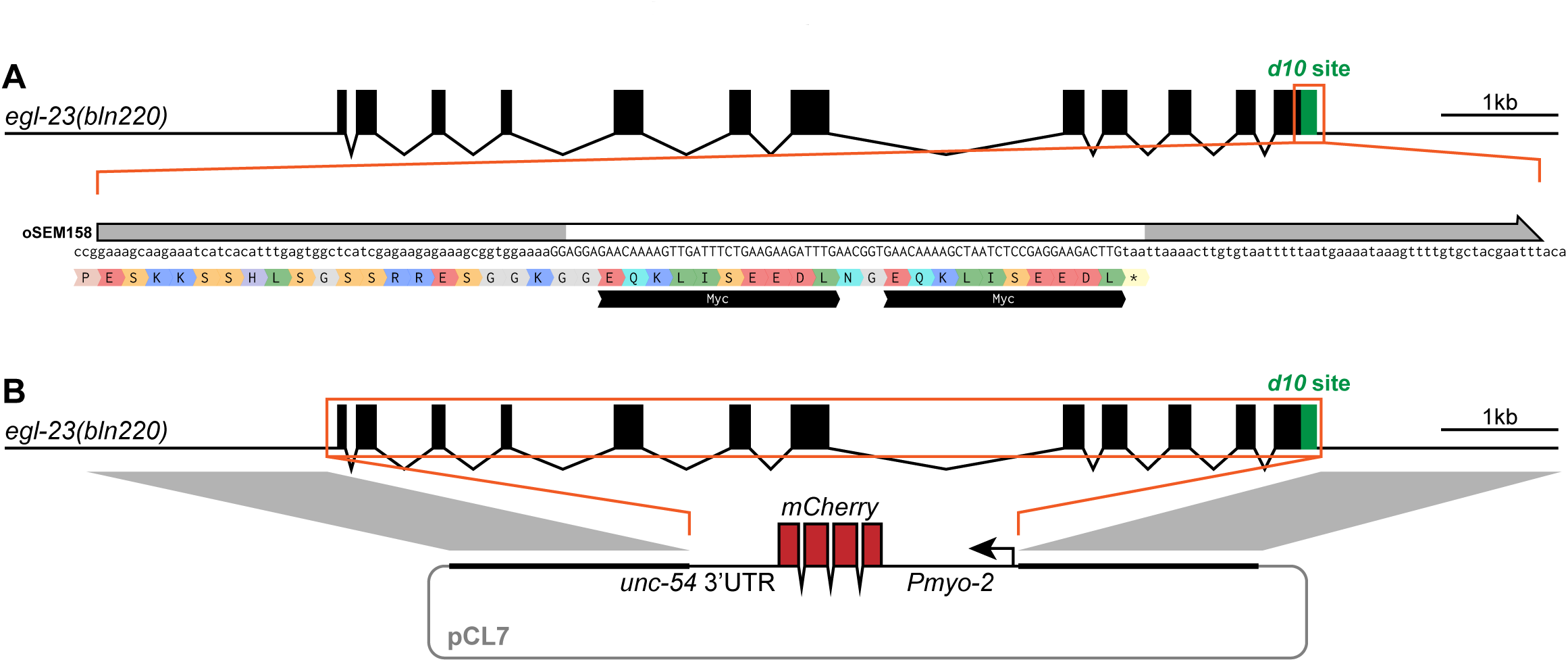
**A** Insertion of a 2xMyc tag into the *egl-23* locus using a single-strand DNA template. Correspondence of homology regions between the ssON repair template and genomic locus is indicated in gray. The sequence of the resulting fusion protein is indicated below the DNA sequence with single-letter amino acid code. Black bars labeled “Myc” indicate the position of the *myc* tag sequences. **B** Deletion and replacement of the *egl-23a* locus by a *Pmyo-2::mCherry* reporter transgene. Correspondence of homology regions between the plasmid repair template (pCL7) and genomic locus is indicated in gray. The *Pmyo-2::mCherry::unc-54 3’UTR* transgene is inserted in the reverse orientation relative to the *egl-23* gene.

**Figure 3 Supplement 1.**
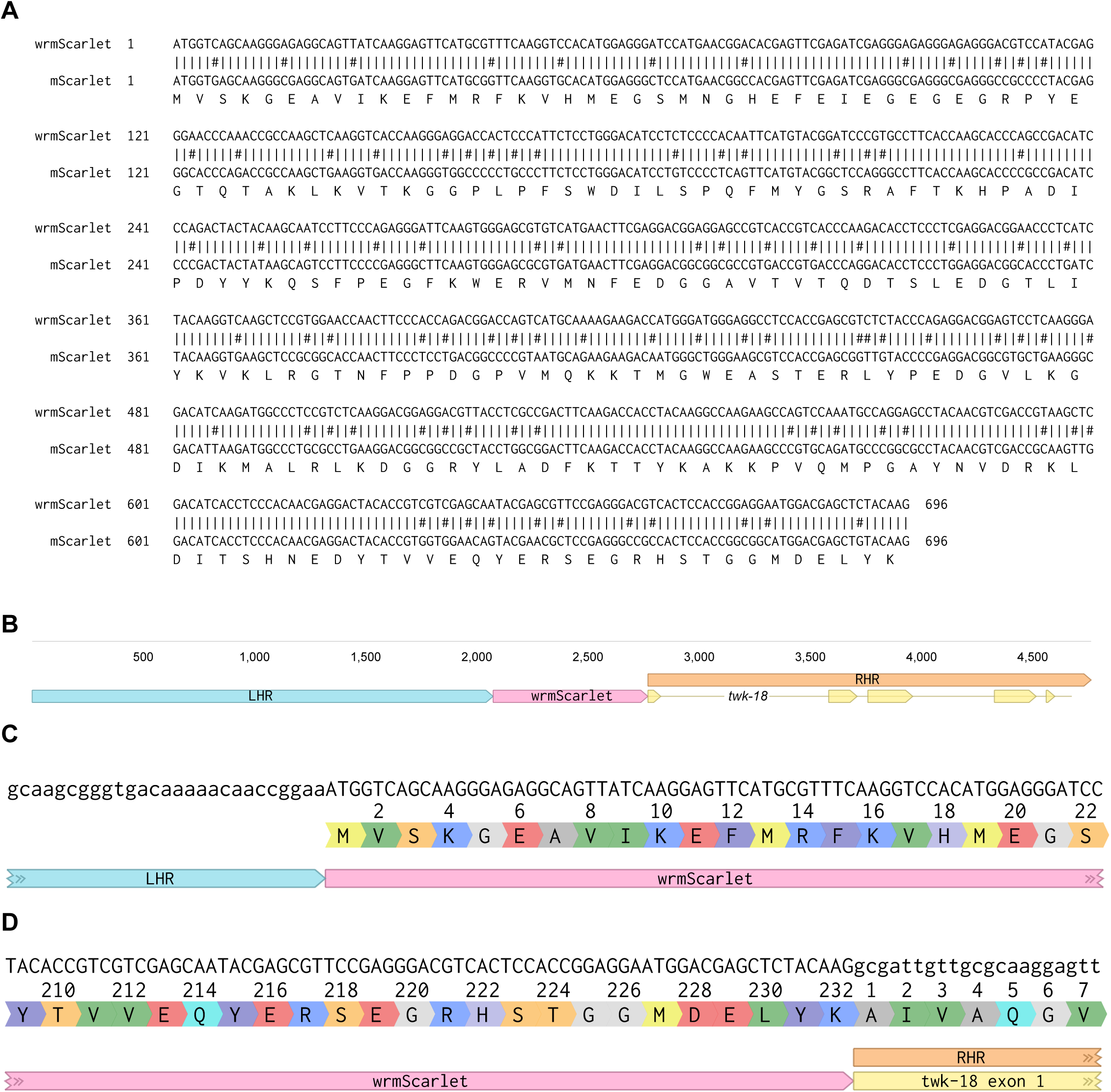
**A** Codon-optimized wrmScarlet vs mScarlet sequence alignement. **B** Schematic representation of the pSEM87 *wrmScarlet::twk-18 repair template*. LHR and RHR indicate left and right homology regions, respectively. The five first exons of *twk-18* present in the RHR are indicated in yellow. Scale bar in base pairs. **C** 5’ junction of wrmScarlet to *twk-18*. The sequence of the resulting fusion protein is indicated below the DNA sequence with single-letter amino acid code and corresponding amino acid positions. **D** 3’ junction of wrmScarlet to *twk-18*. The sequence of the resulting fusion protein is indicated below the DNA sequence with single-letter amino acid code and corresponding amino acid positions.

## Supplementary Tables

**Supplementary Table 1:**
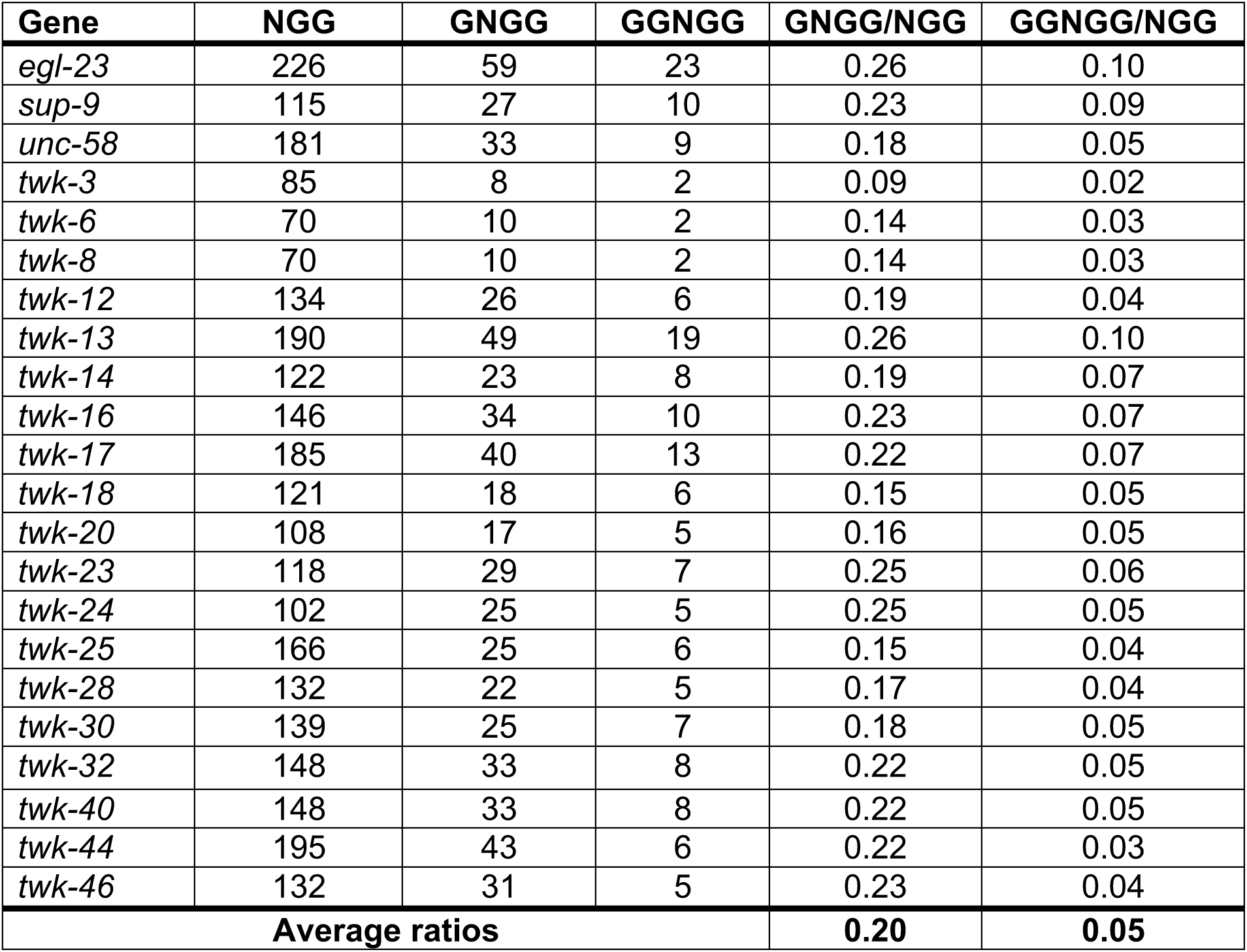
Prevalence of GNGG and GGNGG protospacers in and close to exons of two-pore domain potassium channel genes.

**Supplementary Table 2:**
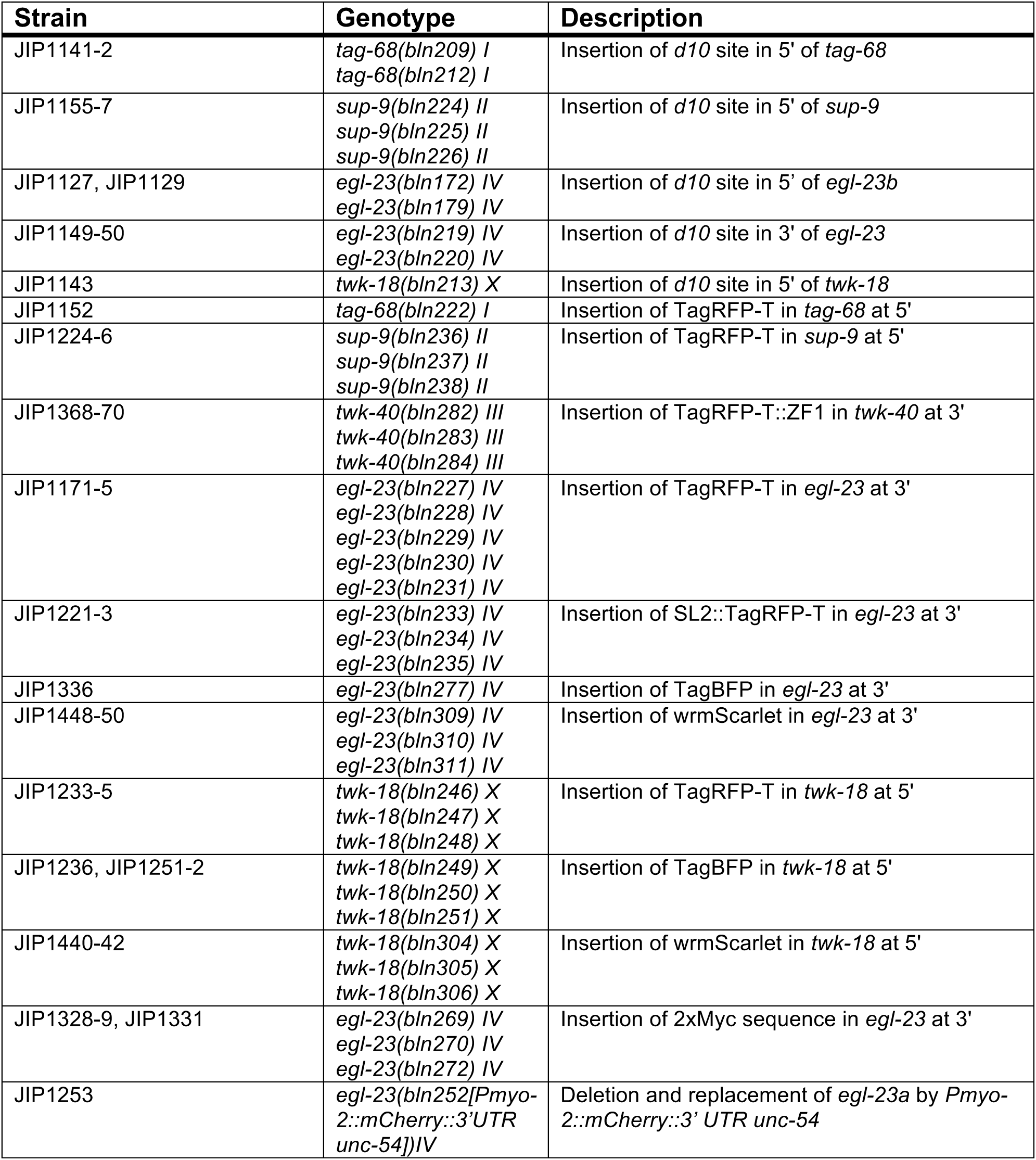
Strain list

**Supplementary Table 3:**
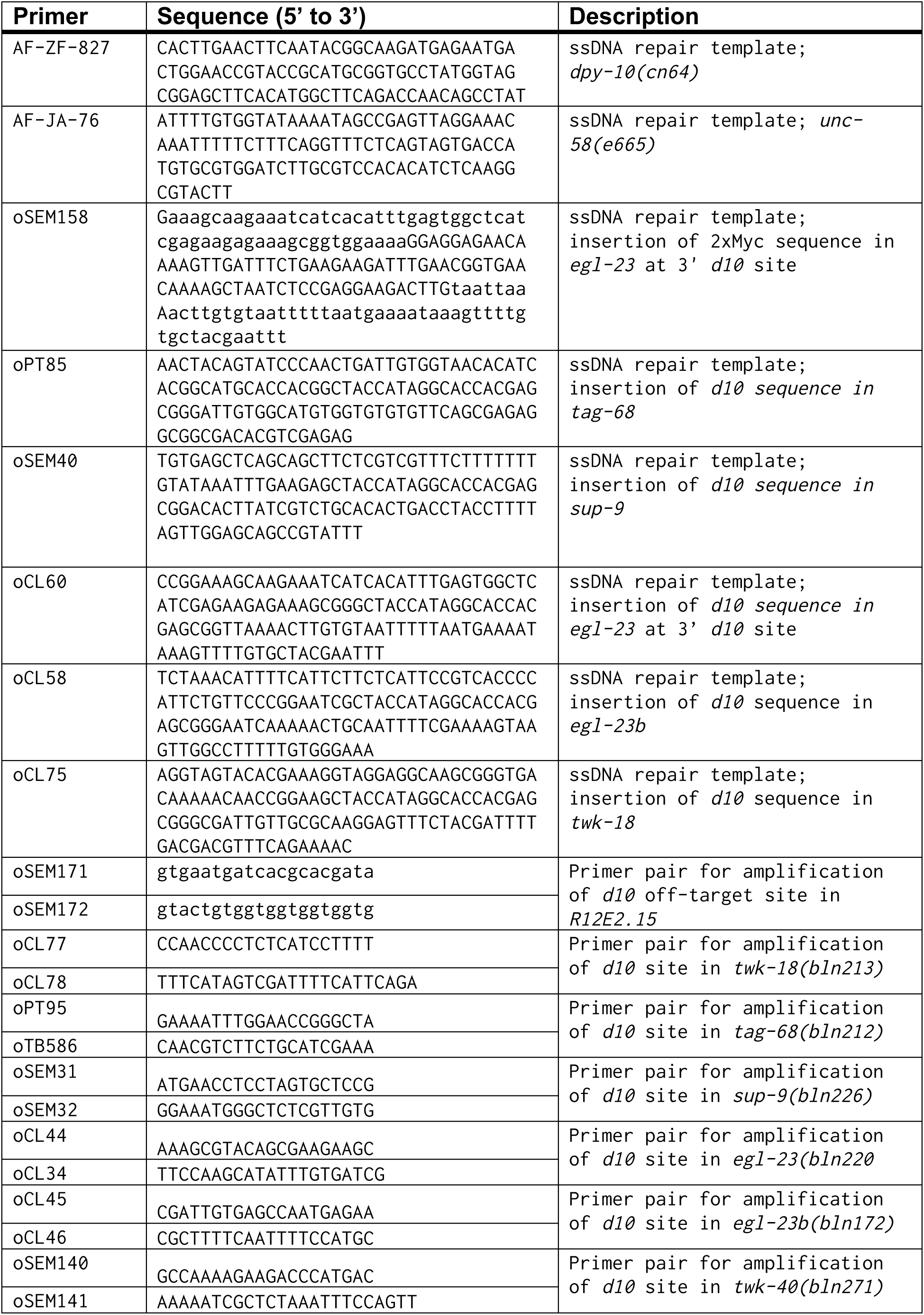
List of single strand oligonucleotides

**Supplementary Table 4:**
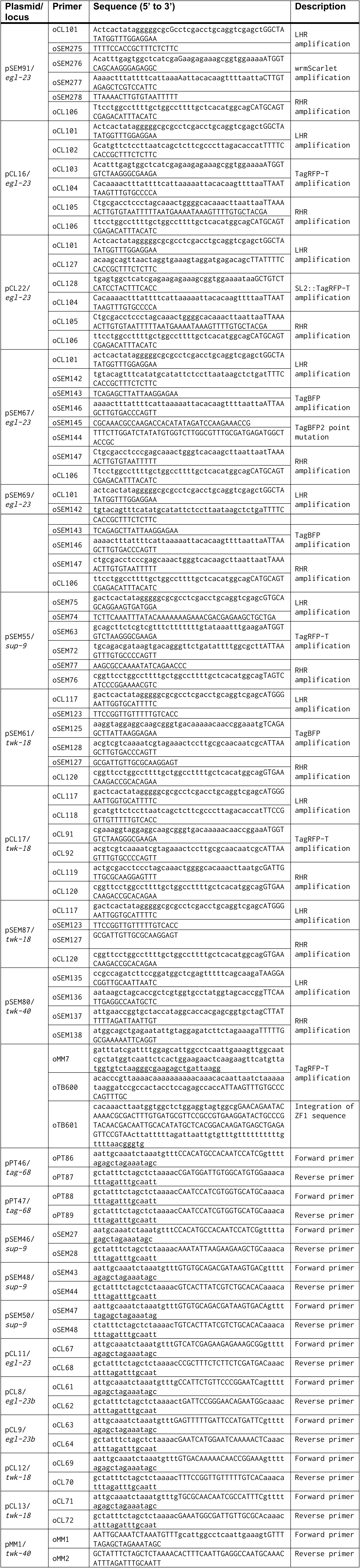
List of plasmids

## Supplementary Methods

### Building sgRNA expression vectors using pPT2

This protocol describes the steps and tools used to generate sgRNA expression vectors using the pPT2 vector backbone. See the materials and methods section for the required reagents (e.g. Gibson assembly reagents, sequencing primers).

We use the excellent online service www.benchling.com for sgRNA, oligo and vector design.

#### I) Identifying a suitable protospacer motif

- A protospacer is a 19-20 bp sequence flanked at its 3’ end by an NGG PAM (protospacer adjacent motif). Different online tools are available to identify possible protospacers in a region of interest (crispr.mit.edu; tefor.net/crispor/crispor.cgi; benchling.com).
- When multiple protospacer sequences are possible, select the closest (to the site to engineer) and/or the most specific sequence (use the off-target prediction tool provided by benchling for example). In general, four non-matching bases should be enough to significantly reduce off-target cutting, especially if the mismatches are located in the 3’ region of the protospacer (Hsu et al. 2013).

#### II) Building the sgRNA vector sequence *in silico*

- The pPT2 vector contains the U6 promoter and 3’ UTR of *K09B11.12* (Friedland et al. 2013) and two restriction sites (PmeI and SexAI) to linearize the vector, followed by the invariant sgRNA scaffold sequence (see Figure 1A).
- To generate the sgRNA expression vector sequence, insert the protospacer sequence (**without** the PAM, i.e. NGG) between the U6 promoter and the sgRNA scaffold as shown in figure 1B.
- If the selected protospacer sequence does not begin with a guanine residue, add this nucleotide manually to the 5’ of the protospacer (i.e. resulting in a “19+1” bp insertion in pPT2 since this protospacer is only 19 bp, see figure 1B).
- Name this vector pXYn where XY are the initials of the person building the vector and n the number of the vector. Accordingly, the protospacer sequence is then labeled CRpXYn (generate a “feature” with the sequence to identify it easily in the genomic sequence).
- Generate one 60 bp oligonucleotide centered on the protospacer sequence as shown in figure 1C (forward or reverse). **Gibson assembly can be performed with a single primer.** Alternatively, generate two complementary 60 bp oligonucleotides centered on the protospacer sequence as shown in figure 1C.

**Figure 1.**
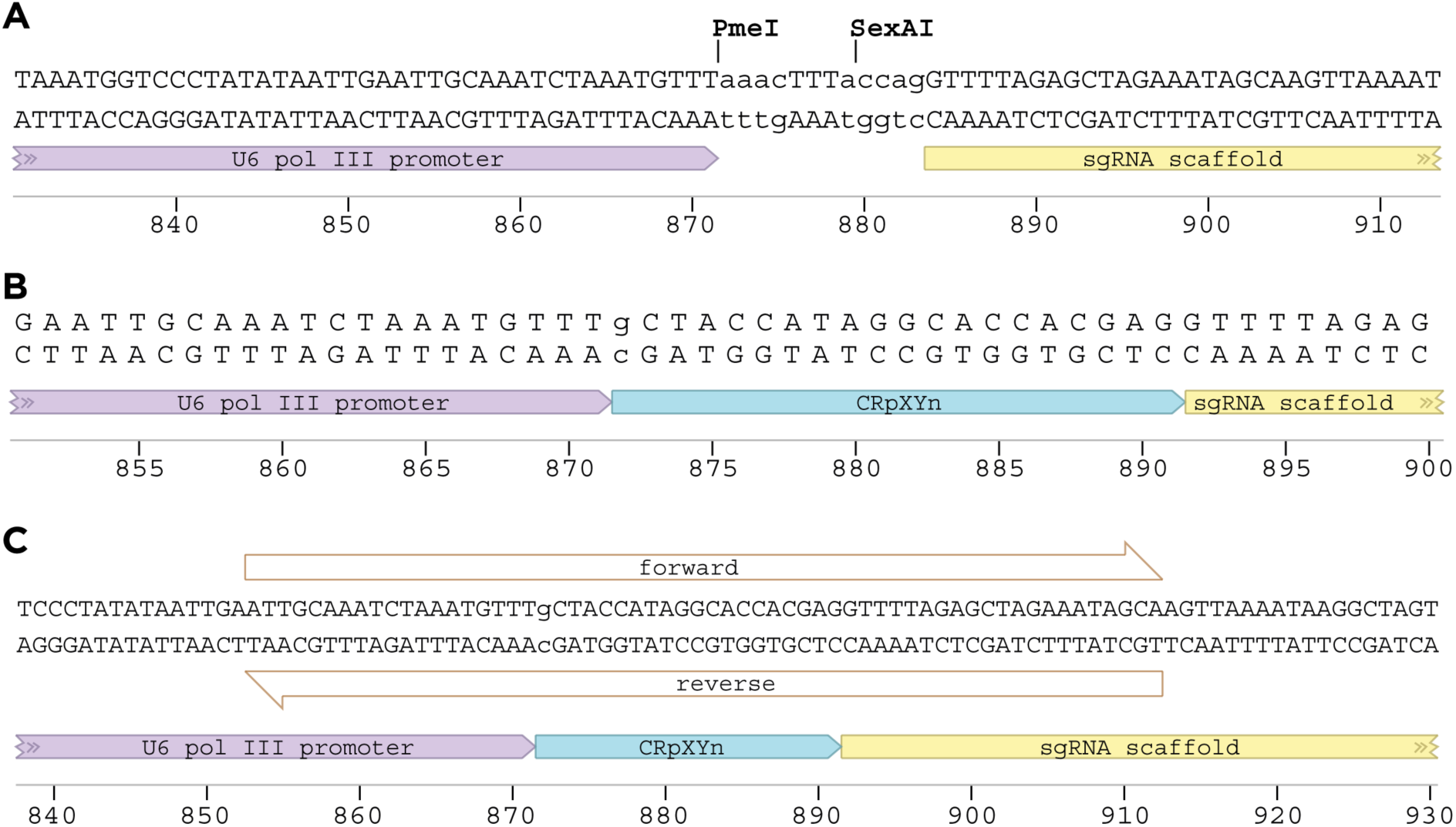
Insertion of a protospacer sequence into the pPT2 sgRNA expression vector. Note that a single primer (forward or reverse) is sufficient to complete the Gibson assembly reaction.

#### III) Building the sgRNA vector

The protospacer sequence is incorporated into the pPT2 vector as follows:

1 | **Gibson assembly**

- Thaw an aliquot of Gibson Master Mix and keep on ice.
- Mix 100 ng of linearized pPT2 vector with 1 μL of 100 μM (or 0.1 nmole) of single strand oligonucleotide and add water up to 5 μL if necessary.
- Add 15 μL of Gibson Master Mix to the DNA mix.
- Incubate at 50°C for 15 to 60 minutes (60 minutes is optimum).
- Transform 5 μL of this reaction, and grow on LB+Ampicilin plates.
- Perform a control experiment (Gibson assembly without primer dimer) for each new batch of linearized pPT2 vector.

2 | **Sequence validation**

- Due to the high efficiency/specificity of Gibson assembly, colony PCR is not required.
- Validate the resulting vector by sequencing with pJET1.2fwd or pJET1.2rev.

3) ***C. elegans* transformation**

Usually, sgRNA vectors are injected at 50 ng/μL.

## Materials and Methods

### pPT2 sequence file

The annotated sequence file for the pPT2 vector can be downloaded as a Genbank format (readable in ApE and benchling.com) at the following link: http://www.excitingworms.eu/resources/pPT2.gb.

### Homemade Gibson Assembly Reagents

Based on *Methods in Enzymology, Volume 498*

*CHAPTER FIFTEEN part 5:Enzymatic Assembly of Overlapping DNA Fragments* (Daniel G. Gibson)

**<u>2 M MgCl_2_</u>** Dilute 10.16 g MgCl_2_ H_2_O for a final volume of 25 mL.

**<u>5X ISO Buffer, 2 mL (store at -20°C)</u>**

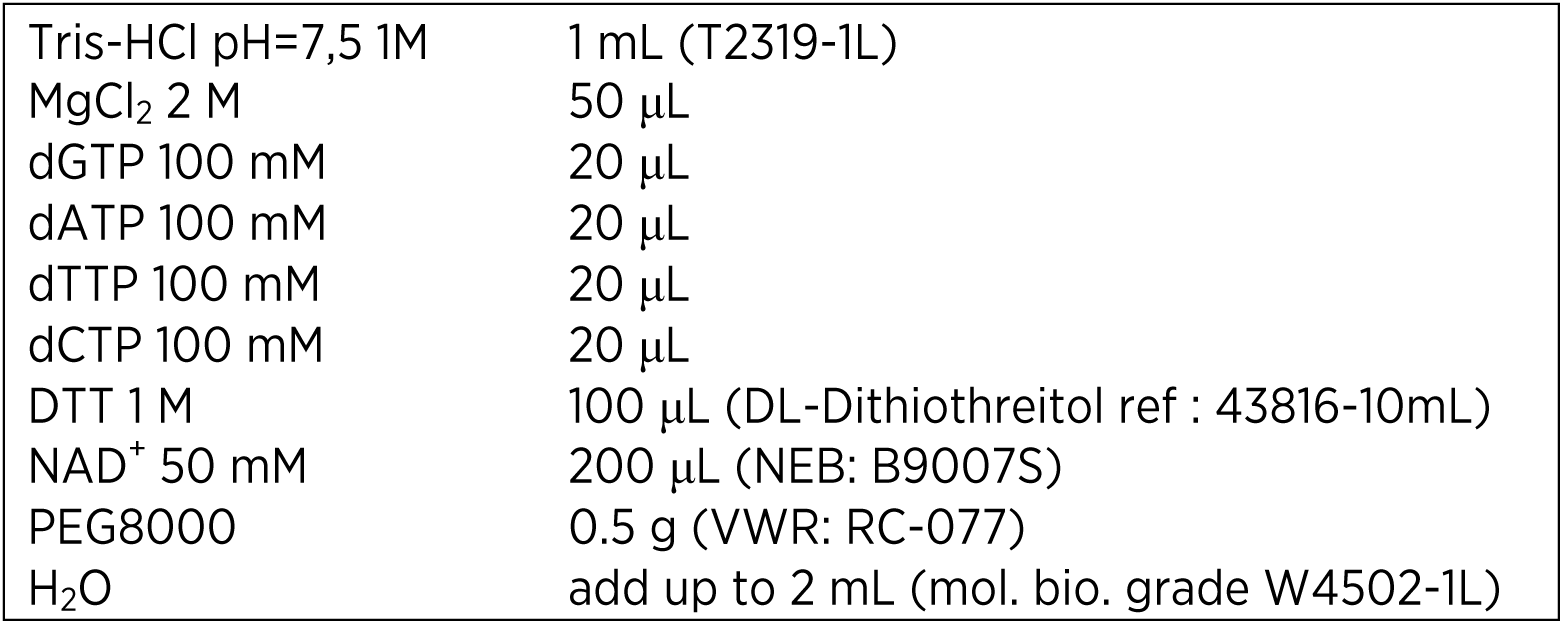

• Prepare 320 μL aliquots and store at -20°C.

**<u>Gibson Master Mix, 1.2 mL (store at -20°C)</u>**

For the assembly of DNA molecules with overlaps of 20-80 bp.

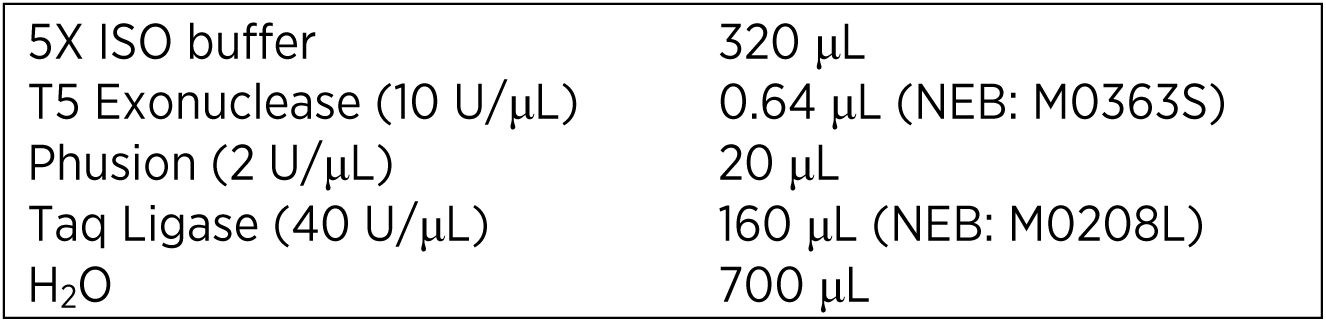

Note: For overlaps that are larger than 80 bp, 3.2 μL exonuclease is used in this mix.

- Separate into 50 μL aliquots. Store at -20°C. The enzyme remains active after 10 cycles of freeze-thaw.

**<u>Sequencing Primers</u>**

pJET1.2fwd 5’-cgactcactatagggagagcggc-3’

pJET1.2rev 5’-aagaacatcgattttccatggcag-3’

### [Alternative protocol used for primer dimer assembly into pPT2] III) Building the sgRNA vector

- The protospacer sequence is incorporated into the pPT2 vector as follows.

1 | Linearize pPT2 using the *PmeI* and *SexAI* restriction enzymes.
2 | Purify the linearized pPT2 vector using your method of choice (we use Qiagen Gel Purification).
3 | **Hybridize oligonucleotides** (using a thermocycler)

- Add 1 μL of each oligonucleotide (at 100 μM) to 18 μL of water.
- Run the program below on a thermal cycler to anneal primers.
- Add 30 μL of water to the resulting sample.

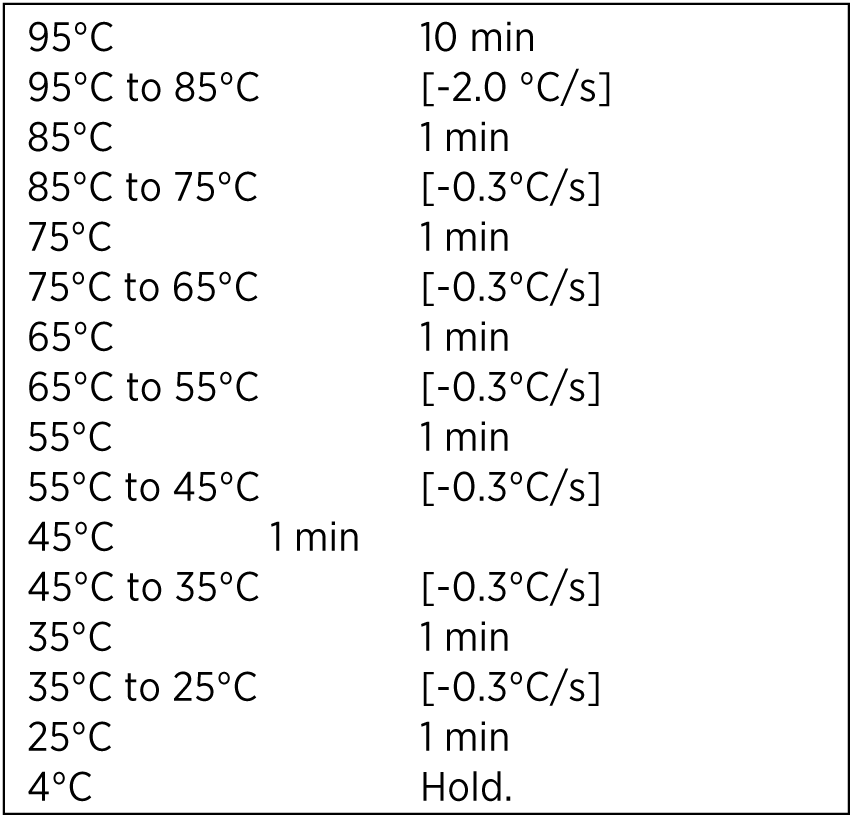

4 | **Gibson assembly**

- Thaw an aliquot of Gibson Master Mix and keep on ice.
- Mix 100 ng of linearized pPT2 vector with 1 μL of hybridized oligonucleotides and add water up to 5 μL if necessary.
- Add 15 μL of Gibson Master Mix to the DNA mix.
- Incubate at 50°C for 15 to 60 minutes (60 minutes is optimum).
- Transform 5 μL of this reaction, and grow on LB+Ampicilin plates.
- Perform a control experiment (Gibson assembly without primer dimer) for each new batch of linearized pPT2 vector.

5 | **Sequence validation**

5) ***C. elegans* transformation**

Usually, sgRNA vectors are injected at 50 ng/μL.

